# An End-to-end Oxford Nanopore Basecaller Using Convolution-augmented Transformer

**DOI:** 10.1101/2020.11.09.374165

**Authors:** Xuan Lv, Zhiguang Chen, Yutong Lu, Yuedong Yang

**Affiliations:** School of Computer Science, National University of Defense Technology, Changsha, China; School of Data and Computer Science, Sun Yat-sen University, Guangzhou, China

**Keywords:** Oxford Nanopore sequencing, Third-party Base-caller, Attention, Dynamic convolution

## Abstract

Oxford Nanopore sequencing is fastly becoming an active field in genomics, and it’s critical to basecall nucleotide sequences from the complex electrical signals. Many efforts have been devoted to developing new basecalling tools over the years. However, the basecalled reads still suffer from a high error rate and slow speed. Here, we developed an open-source basecalling method, CATCaller, by simultaneously capturing global context through Attention and modeling local dependencies through dynamic convolution. The method was shown to consistently outperform the ONT default basecaller Albacore, Guppy, and a recently developed attention-based method SACall in read accuracy. More importantly, our method is fast through a heterogeneously computational model to integrate both CPUs and GPUs. When compared to SACall, the method is nearly 4 times faster on a single GPU, and is highly scalable in parallelization with a further speedup of 3.3 on a four-GPU node.

## I. Introduction

The MinION sequencer developed by the Oxford Nanopore Technologies (ONT) is the first portable DNA sequencing device. It is rapidly becoming a key instrument in genomics[1] because of several advantages: long reads, small size, low cost, real-time analysis, etc. The ONT sequencing is performed as a single-strand DNA passing through a nanopore embedded in a membrane where a voltage difference is applied to generate a small electrical current. The nucleotides present in the nanopore affect the current in a way that can be measured and translated into the corresponding bases[2]. The process of translating the raw electrical signals into DNA sequences is known as basecalling.

Basecalling has a pivotal role in ONT sequencing, largely determining the usability of sequencing results for downstream applications [3–5]. However, generating high-quality sequencing reads remains a challenging task. Despite the capability of the MinION device to produce long reads, the nanopore sequencing reads often show an error rate of 10% or even higher[6], which can be introduced by the noisy and stochastic data in raw electrical signals, as well as the limited accuracy of existing basecalling tools. In the MinION sequencer, each signal value is determined by the multiple nucleotides reside in the nanopore (generally 5 for R9.4 pore), resulting in a large number of base combinations (4^5^=1024) and making it difficult to precisely translate the raw signals into nucleotide sequences. Additionally, a successful MinION run can produce approximately 1.5-2.0 million electrical signals per second[7], magnitude faster than the speed of most existing basecalling tools. In a struggle to keep up with the data generation speed of ONT sequencer, basecallers of fast versions have to sacrifice accuracy performance. Therefore, it is essential to develop a new basecaller that balances accuracy and efficiency.

With the development of deep learning techniques, deep learning has been widely used in many fields[8–10], including the basecalling. ONT has provided several official basecalling tools including Albacore[11], Guppy, Scrappie and Flappie. Albacore runs on CPUs while Guppy performs on GPUs and both of them are only available to ONT’s full members. ONT has suspended the development of Albacore opting for improved performance of Guppy. Scrappie is a technology demonstrator for testing new algorithms which will be the incorporation of new versions of Guppy. Albacore, Guppy, and Scrappie were all developed using an alternating reverse-GRU and GRU architecture. Scrappie has recently been replaced by Flappie, which applies Connectionist Temporal Classification (CTC)[12] to decode bases. Flappie shows higher accuracy than Scrappie but suffers from low speed performance in Wick’s research[13]. Some third-party basecallers have also been released in recent years. Nanocall[14] is an open-source off-line basecaller based on hidden Markov models (HMMs) while incapable of detecting homopolymer repeats[15]. DeepNano[16] predicts the DNA sequences using recurrent neural networks (RNNs), but similar to Nanocall, its application is limited to R7.3 and R9.0 data. BasecRAWller[17], based on unidirectional RNN, provides streaming basecalling but at the expense of accuracy. Nanocall, DeepNano and BasecRAWller are no longer developed. Chiron[18] that combined convolution neural network (CNN) and RNN together with a CTC decoder runs quite slow and thus can only be used on small datasets. More recently, three models, MinCall[6], DeepNano-blitz[7] and SACall[19], were proposed. MinCall used CNN instead of RNN to improve parallelism, however, it was released without well-trained models and trained based on data for R9 chemistry which is currently replaced by R9.4. DeepNano-blitz was developed as a real-time basecaller on CPUs at a cost of accuracy and it is not trainable. SACall, the first basecaller utilizing attention mechanism, still has room for improvement whether in accuracy or speed.

Transformer[20] composed of attention blocks and feed-forward network (FFN) has been widely used in natural language processing and similar tasks[21]. A recent study[19] has demonstrated that it can be successfully applied to the nanopore basecalling task. However, the conventional attention mechanism emphasizes too much on the modeling of local relationships and the FFN layer takes up much of the computation while can hardly perform any contexts captures[22]. To this end, we developed CATCaller based on the Long-Short Range Attention and flattened FFN layer to specialize for efficient global and local feature extraction and increase the capacity of attention without consuming more computation resources. CATCaller is an open-source trainable tool that users can run basecalling directly or re-train it on their own dataset. We tested our model on nine different bacterial genomes and made a comparison with Albacore, Guppy, and the newly released SACall. CATCaller was shown to achieve better performance in terms of read accuracy and error rate. Furthermore, we found that our model runs 13 times faster than its similar method SACall through gpu-based parallel optimization. CATCaller is freely available at https://github.com/biomed-AI/CATCaller.

## II. MATERIALS AND METHODS

### A. Datasets

Although human DNA always attracts much research interest, bacterial genomes are more applicable to the development and examination of ONT basecalling tools. This is due to the fact that bacterial samples allow for a ground truth reference sequence more confident than that provided by complicated eukaryote genomes when measuring accuracy. For this reason, Wick et al.[13] built a training set for Klebsiella pneumoniae using 50 different isolate genomes, including 30 K. pneumoniae, 10 other species of Enterobacteriaceae, and 10 Proteobacteria Species. Besides, 20 reads from each of the 50 genomes were placed into validation set (1000 reads in total), leaving 226116 reads for training. We randomly selected 1/10 reads from 50 genomes to establish our training set. As for test sets, we employed 9 additional genomes provided by Wick et al.[13] as well. These include 3 independent K. pneumoniae genomes (Klebsiella Pneumoniae NUH29, Klebsiella Pneumoniae KSB2 and Klebsiella Pneumoniae INF042) different from the training set and 6 genomes from other bacterial species (Acinetobacter pittii, Haemophilus haemolyticus, Shigella sonnei, Serratia marcescens, Stenotrophomonas maltophilia and Staphylococcus aureus). Accurate reference sequences are also included in their dataset and were generated by hybrid assembling high-quality Illumina reads.

### B. Data preprocessing

The raw signals generated via passing DNA sequences through nanopores can be labeled by ONT default basecalling tools, such as guppy and albacore (https://community.nanoporetech.com). However, the basecalled reads show a fairly high error rate. To get accurate label sequences and increase the quality and usability of our training data, we followed the pre-processing steps used by Huang et al.[19] and Wick et al.[13] The preprocessing procedure before training can be summarized as follows:

1. Trim each raw read at the fast5 level by removing low-variance open-pore signals from the start and end of the signal sequence.
2. Translate the trimmed reads using Guppy basecaller or other existing basecalling tools.
3. Align the basecalled reads to the corresponding reference by minimap2[23] and filter the reads once more according to the quality of the signal-to-reference alignment.
4. Utilize the re-squiggle module of Tombo (https://github.com/nanoporetech/tombo) to rectify mismatched bases and then extract target base sequences from re-squiggled fast5 files.

It should be noted that these preprocessing steps are only necessary for training, and not used in the test.

### C. Model architecture

In the MinION sequencer, a nanopore current measurement is associated with multiple nucleotides since there are several bases present in the nanopore when a current change is detected. An event is defined as a period of raw signals corresponding to a particular DNA context. Consecutive events correspond DNA contexts should differ by one base but in practice they may be the same or differ by more than one bases due to the noisy data during sequencing. Therefore, it is not a trivial work to capture the complicated correlations between each signal and its surroundings. Here, we employed a convolution-augmented transformer architecture instead of opting for the already adopted RNN[7, 16] or CNN[6] approaches. As shown in Fig.1, the CATCaller model has two normal convolution layers and N encoder blocks followed by a fully-connected layer to produce the predicted probabilities of each base. At the top of the architecture is a CTC[12] decoder to output final DNA sequence. Each encoder block is composed of layer normalization, a two-branch long-short range extractor and a flattened FFN block. In this work, we set parameter N = 6.

**Fig. 1.**
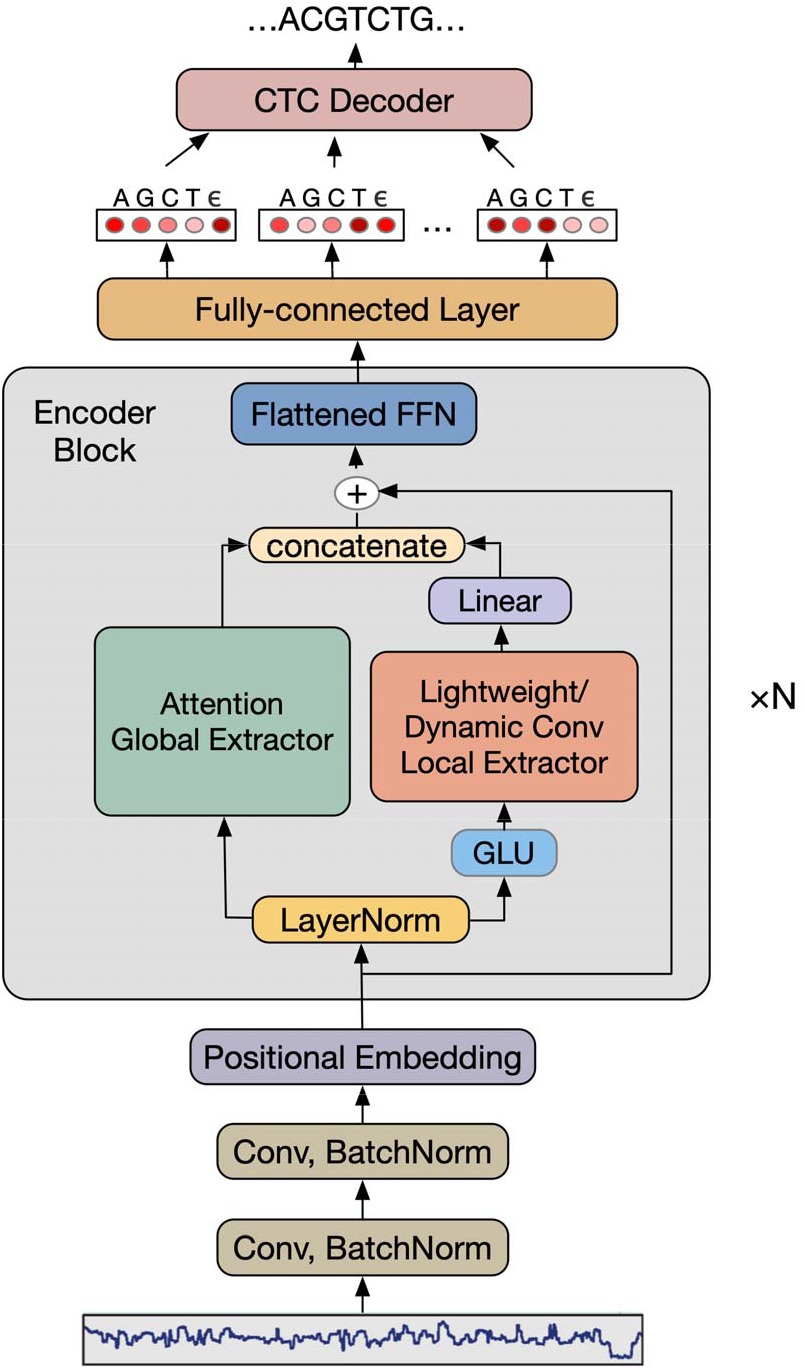
Schematics of CATCaller model, where Conv is the abbreviation for convolution layer and Linear stands for a fully connected layer.

#### Normalization and re-segmentation

Most existing basecallers obtain normalized signals by calculating the mean of data acquisition values over basecalled events, picoamp values and median values in steps. Here, we applied a median absolute deviation (MAD) scale method which directly derives median values from raw electrical signals.

This straightforward normalization is much simpler than multistep procedures but was demonstrated to achieve a comparable performance[24]. The normalized signal data can be calculated as

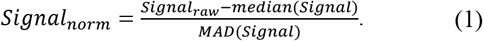

After normalization, we re-segmented signal sequences and the mapped reference bases into chunks by sliding a 2048-length signal window and a 300-length label window. Then, the vectors of preprocessed signal and label segments were fed into the neural network.

#### Sequence embedding

The MinION sequencer can produce ultra-long reads up to 882kb[6], which brings much computational pressure to the encoder block, especially for the attention module. To down sample signals and reduce the computational costs, we employed a convolution module just like SACall[19] model used before the embedding layer. The convolution module contains two 1D convolution blocks, each of which consists of a normal 1D convolution layer followed by a batch normalization layer and a rectified linear unit (ReLU) function[25]. The channel dimension of the input signals was firstly increased from 1 to *d_model_/*2 in the first convolution layer and then doubled to *d_model_* in the second one. Both of the two convolution filters have a kernel = 3, padding = 1 and stride = 2. The length of each input signal sequence shrinks to 1/4 of the initial value after the downsample block.

Before the encoder block, we adopted the sinusoidal positional encoding method proposed by Vaswani et al.[20] to remember the sequential information of the signals in a long read.

#### Long-short Range Attention

As shown in Fig.1, we replaced the normal attention module with the *Long-Short Range Attention* (LSRA) inspired by Wu et al.[22]. Although the attention mechanism has been confirmed successful in various tasks, it still suffers from limitations of high computational requirements and the redundancy of local relationship modeling[26, 27]. To address these problems, LSRA splits the normal multihead attention block into two branches along the channel dimension. The left branch maintains the conventional attention module with a channel dimension of *d_model_/*2, while the right part processes the other half of channels by using lightweight or dynamic convolution module.

For the long-distance relationship modeling, we used the naive multi-head self-attention architecture from the Vaswani transformer[20]. The multi-head self-attention module constitutes *ℎ* heads performing scaled-dot attention mechanism in parallel. Each scaled-dot attention layer takes signal embedding as input and firstly computes three matrices: the queries Q, the keys K, and the values V. These matrices are obtained by applying three linear projections to the input signal embedding. These operations used in the multi-head unit are as follows:

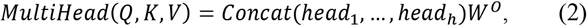

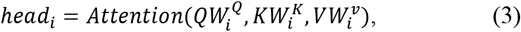

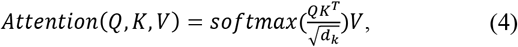

where 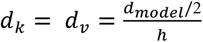 and *i* denotes head (*i* = 1,2, …, *h*).

Another group of heads specializes in the local feature extraction by using a dynamic convolution module introduced by Wu et al[28]. They found that both lightweight convolution and dynamic convolution can achieve results of comparable accuracy to the self-attention but are simpler and more efficient. Lightweight convolution based on the depthwise separable convolution[29] shares weights in each group of channels and the weights are normalized along the temporal dimension using a softmax function. The lightweight convolution is calculated as follows:

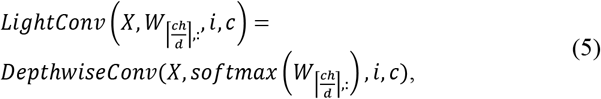

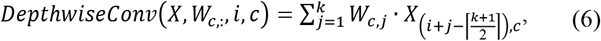

where *d = d_model_/*2, *i* stands for the *i^th^* element in the sequence, *k* is the kernel width, *c* corresponds to the output channel, *h* represents the group number of channel equaling to the head number in the attention module. Dynamic convolution built on the lightweight convolution dynamically learns a new kernel at every time step by using a function *f*: ℝ^*d*^ → ℝ^*h*×*k*^:

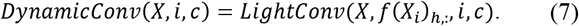

We deployed a Gated Linear Units (GLU) [30] and a fully-connected layer before and after the convolution module respectively and the kernel sizes are [3,5,7,31×3] for the overall six encoder blocks.

In the original design of the Vaswani transformer[20], the FFN firstly increases the input channel dimension from *d_model_* to *d_ff_* (usually *d_ff_* is 4× larger than *d_model_*) and then reduces it to *d_model_* through two linear layers. Wu et al.[22] demonstrated that the computation is dominated by the FFN while FFN can hardly perform any contexts captures. Therefore, we replaced the original FFN with the flattened FFN in which both the channel dimensions of input and output are equal to *d_model_*. In this manner, the capacity of long-short range feature extractor was enhanced with a large dimension *d_model_ =* 512 without additionally leading to a higher computation cost than the original transformer.

### D. CTC decoder

After the encoder module is a fully connected layer followed by a log-softmax function to convert hidden states *H* at position *i* to log probabilities:

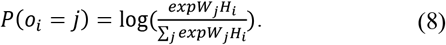

The output *o_i_* predicts the corresponding symbol *j ∈ {A, G, C, T, E}*, where ‘*E*’ stands for a blank symbol. The output is then fed into a CTC decoder based on a beam search algorithm with a beam width *w =* 3. The beam search decoder maintains a set of *w* most probable prefix sequences up to position *i*. The new set of prefixes at position *i +* 1 is generated from the previous set by extending each prefix with all possible character and keeping the top *w* candidates. Notably, each candidate prefix is stored after firstly merging adjacent duplicates into one character and then removing the ‘*ϵ*’ symbols. The algorithm accumulates the scores for each prefix in the beam at each iterate of the search process and in the end selects the one with the highest score as the final output sequence.

### E. Training settings and calling

During the training procedure, we calculated the CTC loss between basecalled and target sequences. An Adam[31] optimizer with a learning rate *lr = 1e*^−4^ was used to minimize the CTC loss. To avoid causing instability of the deep neural network, we used a warmup learning rate scheme to gradually increase learning rate:

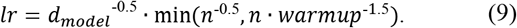

where n is the step number and warmup *=* 10^4^. We used a dropout probability of 0.1 for all dropout layers in our model. CAT-Caller is an open-source tool that users can train their own model with a certain dataset. To take full advantage of the advanced deep learning hardware like NVIDIA’s Volta and Turing GPUs and improve training efficiency, we provide a parallel training interface that supports training on multi-GPUs and multi-nodes with mixed precision using NVIDIA’s APEX library[32]. The batch size for individual GPU is 32.

As an end-to-end basecaller, users can also run basecalling directly with our well-trained model. In addition to keeping the data and model in float32 during prediction, the calling module also supports a ‘half’ mode that uses the float16 for a faster calling speed without sacrificing accuracy. To further reduce the basecalling time and ensure scalibility, we extended our implementation to utilize the immense computational potential of multi-GPU systems through data parallelism and the overlap of encoding and decoding phases.

## III. RESULTS AND DISCUSSION

### A. Evaluation metrics

We adopted nine different bacterial genomes from Wick et al[13] to estimate the performance of CATCaller. None of these benchmark sets was used for training or validation.

We evaluated the basecalled results at the read level using “read accuracy” which measures the identity of each basecalled read relative to the reference sequence. Read identity of each read is defined as

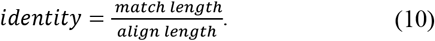

The basecalled sequences were firstly aligned to the corresponding reference using minimap2[23] and then the overall read accuracy was obtained by calculating the median identity of all reads in a sample set. The error rate is adopted as another criterion which is computed as

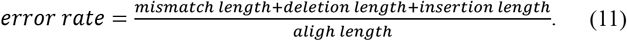

### B. Performance comparison

We compared our method with three other end-to-end basecallers that support R9.4 chemistry, including the ONT official tools Albacore(v2.3.4) and Guppy(v3.2.2) together with the third-party basecaller SACall[19]. We also examined the Guppy-Kp (v2.2.3) model that was trained on the same data as CATCaller. The Guppy was tested using the HAC model (HAC means high accuracy). TABLE I displays the read accuracy of these basecallers on nine benchmark bacterial genomes. CAT-Caller generated the read accuracies of over 90.7% for all test sets and consistently outperformed SACall, Albacore, and Guppy-KP. Guppy performed better on the Acinetobacter pittii and Staphylococcus aureus, especially the latter, which showed a much higher accuracy than on other bacterial samples. This divergence might be caused by the tendency of Guppy’s training set since the accuracy distribution of Guppy-KP was similar to our method. We also calculated the average result of the nine benchmark sets and CATCaller produced the highest accuracy of 91.52%, which was superior to that generated by Guppy (90.92%). The average accuracy of our method is better than SACall, the state-of-the-art basecaller, by 0.4%, suggesting that our enhanced feature extractor is useful. Fig. 2 illustrates the error rates of the above basecallers, among which CATCaller further confirmed its generalization ability by yielding the lowest error rates on most of the bacterial samples.

**Fig. 2.**
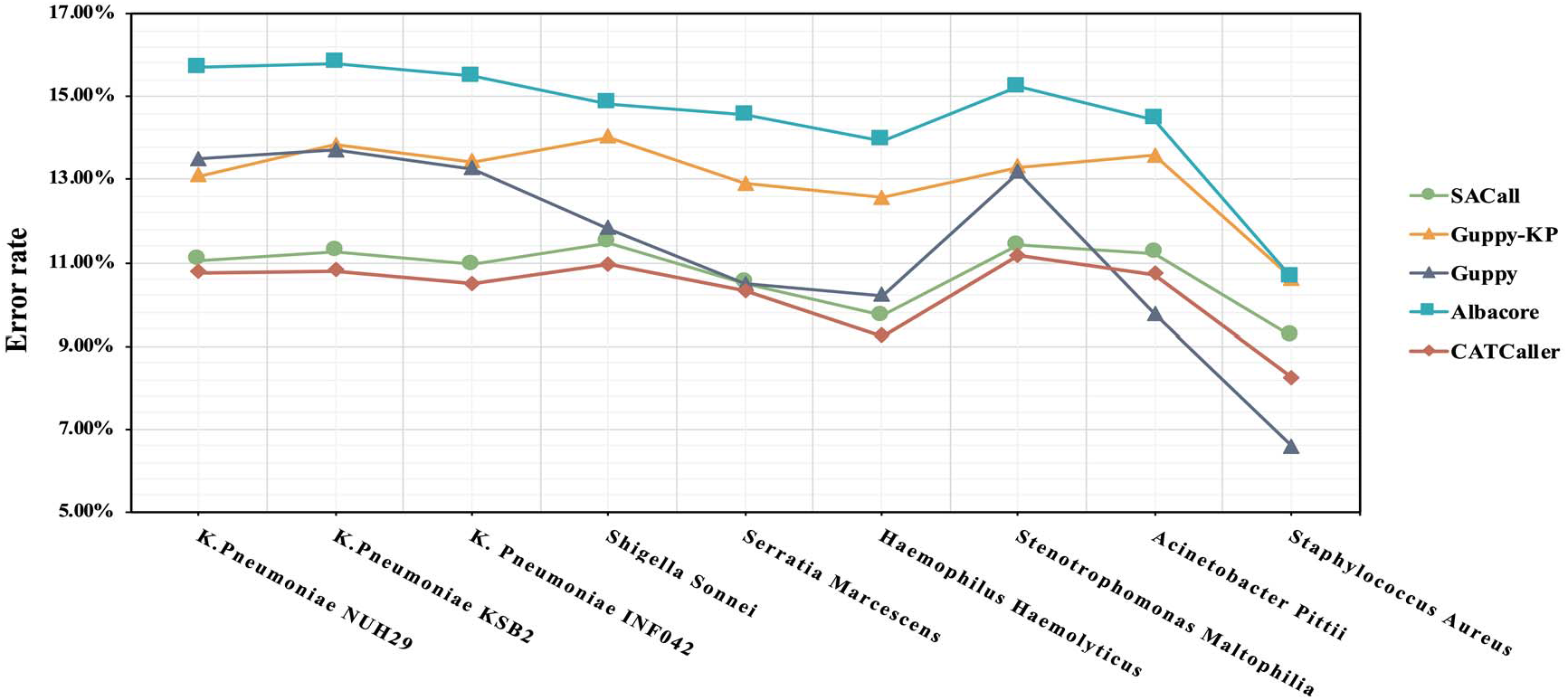
The error rate of different basecallers on nine bacterial genomes

**TABLE I.**
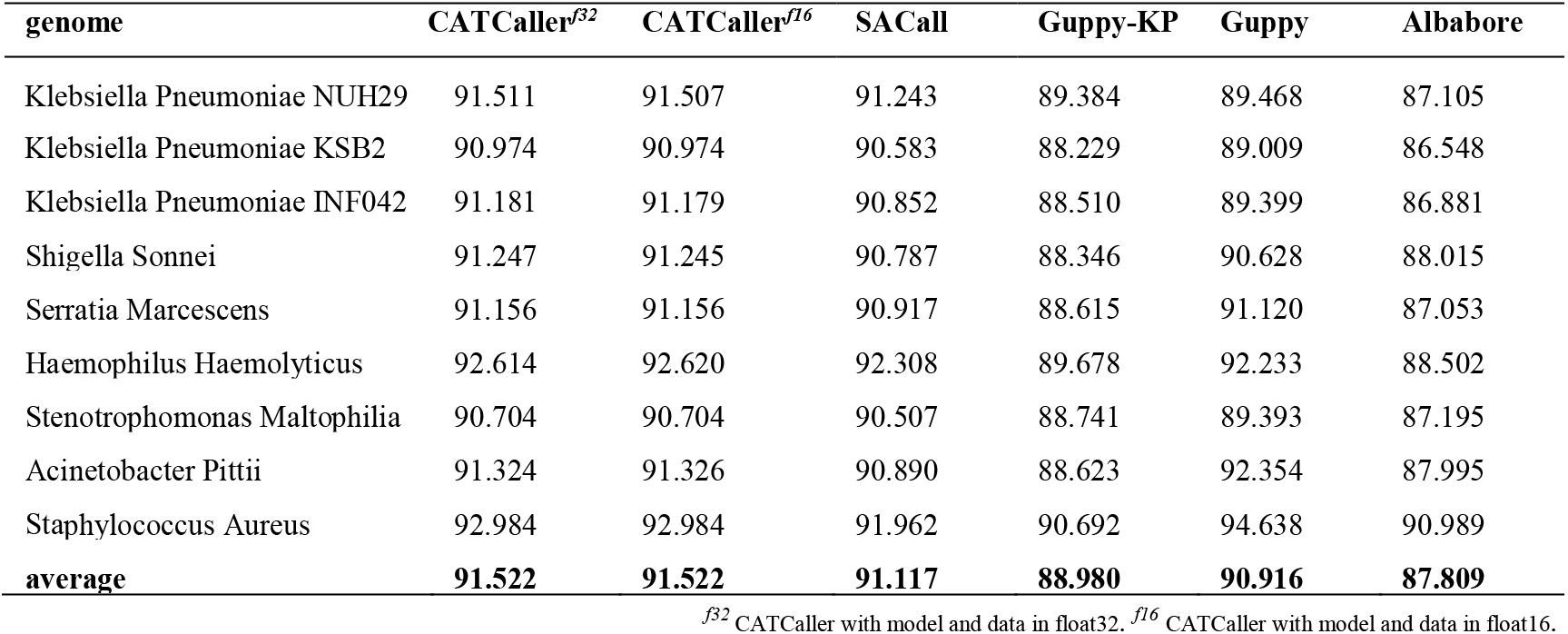
THE READ ACCURACY ON NINE BENCHMARK BACTERIAL GENOMES

As an upgrade of the traditional attention mechanism, the CATCaller model was further compared with SACall in terms of speed performance. TABLE II outlines the average basecalling speed of SACall and CATCaller over nine test sets. We tested the two basecallers on Intel(R) Xeon(R) Gold 6132 CPU @ 2.60GHz with NVIDIA Tesla V100 GPUs. Combining the information from TABLE I and TABLE II, we found that CAT-Caller can be accelerated with little accuracy loss under half precision mode and obtained a 4× speedup over SACall. When scaled to 4 GPUs, CATCaller achieved a speedup by factors up to 13.25 in comparison to SACall. These results suggested that our method, based on convolution-augmented transformer, is able to provide high-quality basecalled reads with a greater speed.

**TABLE II.**
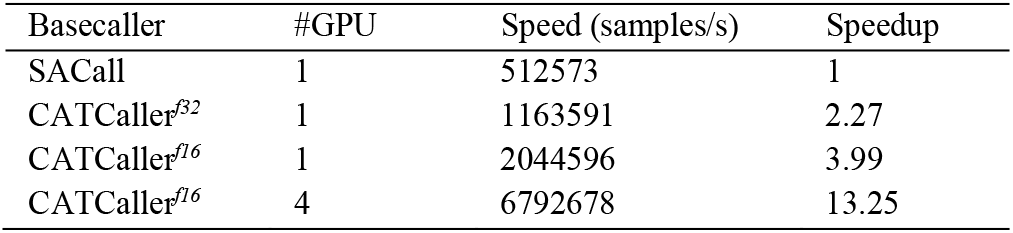
Basecalling speeds measured in samples per second

## IV. CONCLUSIONS AND FUTURE WORK

In this paper, we proposed a new convolution-augmented transformer model, CATCaller, for the nanopore basecalling task. More clearly, we used a two-branch long-short range attention architecture, where one branch specializes in the short-distance dependencies modeling by convolution while another branch focuses on the long-distance relationships with attention module. The LSRA design also introduced a flattened FFN layer that trades off the computation for enhanced feature capturing capability with wider attention blocks. We examined our method on 9 different bacterial samples made by wick et al.[13] and performed a comparison with three other available basecallers. The results showed that CATCaller can achieve better overall performance than RNN based approaches or conventional attention based method. Despite the higher read accuracy of Guppy (v3.2.2) on Acinetobacter pittii and Staphylococcus aureus, CATCaller produced more stable and balanced results. Furthermore, our method shows a speedup of nearly 4× over SACall using a single GPU and the speed can be further accelerated by factors up to 13.25 on a 4-GPU node.

CATCaller can be used as a promising basecaller producing reliable reads or served as a re-basecall tool to further improve the reads quality. This method was developed based on the data sequenced using currently available MinION R9.4 pore, but as an open-source tool it can be easily adjusted for the data sequenced by newer pore types and we believe that it would be useful to the development community.

Current basecallers based on neural networks generally use real data for supervised training and thus the performance may be influenced by the existence of base modifications in their training set. Recent studies[24, 33] have demonstrated that DNA modifications could be detected by analyzing electrical signals during MinION sequencing while attention-based models were rarely used to identify base modifications, which will be included in our future work.

## Acknowledgment

This work was supported by the National Natural Science Foundation of China (61772566, 81801132, and U1611261), Guangdong Key Field R&D Plan (2019B020228001 and 2018B010109006) and Introducing Innovative and Entrepreneurial Teams (2016ZT06D211). We appreciate the availability of the Guppy and Albacore packages, and Dr. Huang Neng from the Central South University for answering questions about SACall.

